# Age-Dependent Fibroblast Programs Govern Regenerative and Fibrotic Tendon Repair

**DOI:** 10.64898/2026.07.22.739917

**Authors:** Ahmet Engin Pazarceviren, Minoo Bastani, Dean Roell, Julia Sheyn, Dave Huang, Oksana Shelest, Mia Orr, Lea Zila, Melodie Metzger, Wafa Tawackoli, Dmitriy Sheyn

## Abstract

Tendon injuries often result in fibrosis, compromising function and predisposing to re-injury. Here, we used a full-width, non-repair Achilles tendon transection model in young (3 weeks old) and adult (18-20 weeks) rats to elucidate the cellular mechanisms governing regenerative versus fibrotic healing. Functional and biomechanical analyses revealed that young tendons recovered motion and load-bearing capacity more rapidly but exhibited more fibrotic early healing. Single-nuclei RNA sequencing identified seven major cell populations within the connective tissue compartment. Adult tendons maintained a “synthetic fibroblast” population marked by upregulated ECM synthesis, reduced stress-related gene expression, whereas young tendons favored expansion of Cxcl12⁺/Lrp6⁺/Gas6⁺ fibrotic fibroblasts linked to oxidative and pro-angiogenic signaling. The young group showed sustained activation of Nox4–Gas6 pathways driving a self-reinforcing fibrotic circuit. These findings define a fibroblast lineage bifurcation that dictates oxidative stress signaling as a key regulator of fibrotic remodeling, highlighting potential therapeutic targets to promote regenerative tendon repair.

## Introduction

Tendon trauma constitutes over half of all musculoskeletal injuries in the United States and affects approximately 10% of the population by the age of 45, including nearly one million athletes worldwide. The socioeconomic burden exceeds $40 billion annually in medical treatment and rehabilitation costs^1^. Despite surgical advances, re-injury rates remain high by exceeding 20% in repaired tendons and this is largely due to the fibrotic nature of adult tendon healing, which leads to persistent scar formation and compromised mechanical function^2^. Tendon fibrosis therefore represents a major barrier to successful regeneration and long-term functional restoration. Adult tendon injuries account for up to 50% of all musculoskeletal injuries, with this number continuing to rise as the population ages and recreational athletic participation increases^3^. Without surgical intervention, functional restoration after tendon rupture is further complicated by fibrotic repair and matrix disorganization^4^. This fibrotic outcome compromises mechanical strength and elasticity, preventing full restoration of function.

Adult tendons exhibit a predominance of fibrotic healing characterized by persistent vascularization and unresolved inflammation, leading to chronic scarring^5^. In contrast, fetal and neonatal tendons in mice display a regenerative phenotype marked by minimal fibrosis, organized collagen alignment, and faster structural recovery^6^. These age-dependent disparities are largely attributed to the abundance and plasticity of tendon progenitor cells and a transient immune milieu that supports tissue restoration rather than chronic inflammation^7^. To study mechanisms of tendon injury and healing, several experimental models have been developed including full-thickness and partial Achilles tendon tear and surgical repair utilizing novel lineage tracing models^8,9^. On the other hand, non-repair models enable assessment of spontaneous tendon healing without repair-associated stabilization and intervention effects that can alter healing outcomes.^10^ Accordingly, we used a non-repair complete Achilles transection to compare endogenous natural regenerative versus fibrotic transcriptional programs at transcriptomics level under a standardized injury condition.

Recent advances in single-cell transcriptomics have revealed substantial cellular heterogeneity within tendon tissue, uncovering specialized fibroblast and progenitor populations that orchestrate healing and matrix remodeling^11,12^. However, single-cell investigations of non-repair tendon healing aiming to elaborate connective tissue clusters remain scarce. Deciphering the intrinsic cellular programs that distinguish regenerative from fibrotic outcomes is therefore essential for understanding the molecular determinants of tendon regeneration and for developing targeted antifibrotic and regenerative therapies.

In this study, adult and young rats were used to examine intrinsic healing following a complete, full-width Achilles tendon transection without surgical repair. This model allowed observing the differences of tenocyte and fibroblast transcriptional states at the connective tissue compartment in natural healing process. We hypothesized that this mode of healing would reveal cellular programs that are amplified during the repair, thus providing an enhanced view of the cell types occurring at the defect zone. These insights highlight a previously underappreciated relationship between progenitor cells, fibroblasts, and the balance between regeneration and fibrosis in tendon healing.

## Results

Age-related changes in tendon recovery and fibrosis morphology. The full width Achilles resection model was designed to evaluate the role of progenitor cell differences in tendon regeneration (Fig. 1A). The functional recovery of the tendons over an 8-week period post-injury was characterized using biweekly gait analysis as previously reported^13^, to understand the locomotor recovery, and biomechanical testing and histological examination to provide an integrated overview of the healing conditions (Fig. 1B).

**Figure 1.**
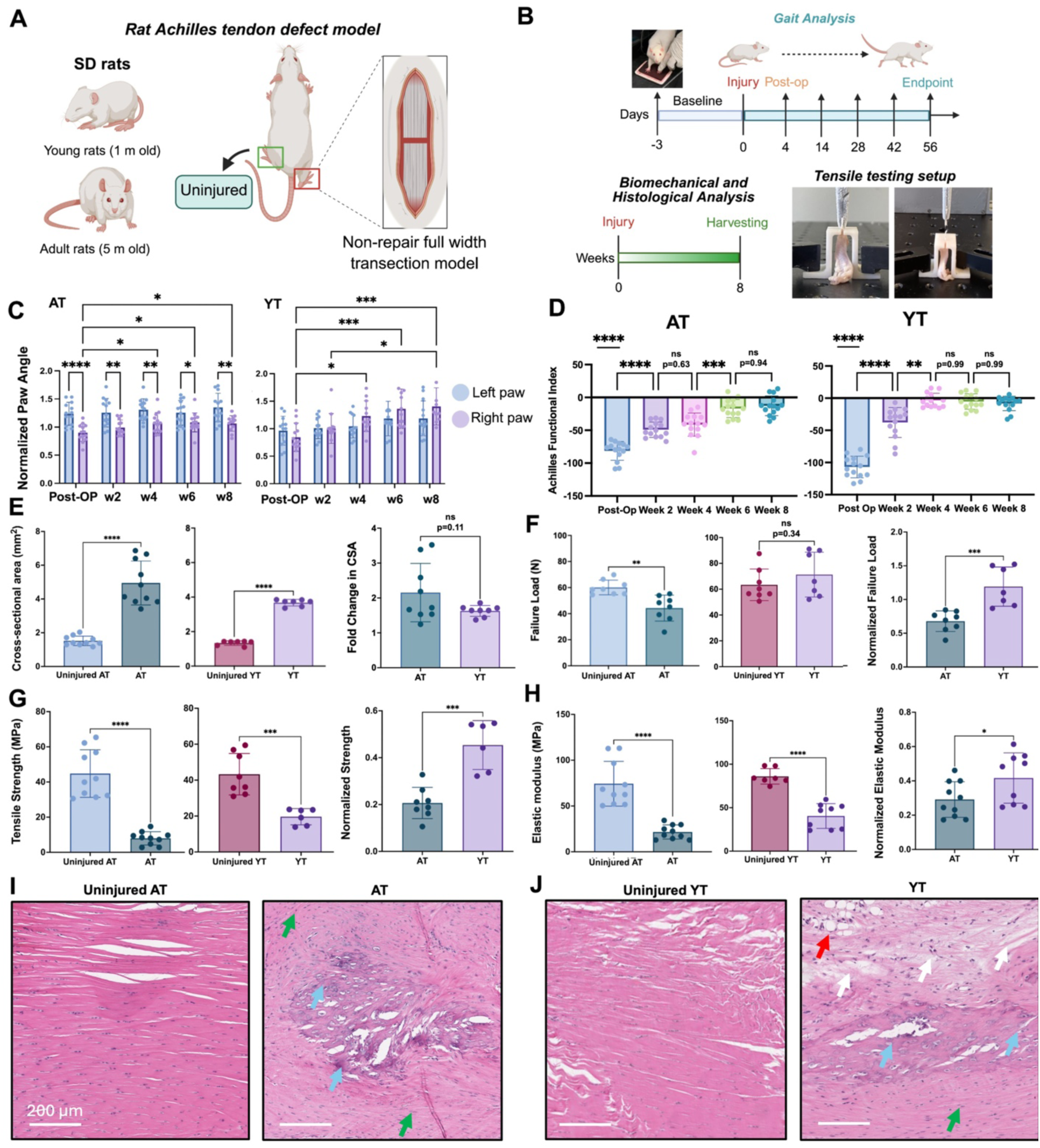
Functional, biomechanical, and histological evaluation of tendon healing in adult and young rats**. A)** Schematic illustration of the experimental design showing full-width Achilles tendon resection in adult (18-20 weeks, AT) and young (3 weeks, YT) Sprague Dawley rats. **B)** Timeline of gait, biomechanical, and histological assessments performed over the 8-week healing period. **C)** Normalized paw-angle analysis showing consistently reduced performance of injured right paws in AT compared with the contralateral uninjured side. YT exhibited rapid recovery by week 2, with right-paw angles exceeding baseline, indicative of compensatory outward stepping (n=14). **D)** Achilles Functional Index (AFI) revealed significantly improved recovery in YT by week 4, while AT displayed slower but progressive improvement, reaching partial recovery by week 6 (n=14). **E–H)** Biomechanical parameters such as cross-sectional area (CSA), failure load, tensile strength, and elastic modulus demonstrated a faster and mechanically superior recovery in YT relative to uninjured AT tendons (n=9). **I)** Representative histological analysis of AT tendons showed restoration of aligned collagenous matrix (green arrows) with a persistent fibrotic scar characterized by disorganized fibers (blue arrows). **J)** YT tendons contained regions of aligned collagen intermixed with immature, non-aligned extracellular matrix, extensive neovascularization (red arrows), and focal tissue discontinuities (white arrows). Statistical significance: *p < 0.05, **p < 0.01, ***p < 0.001, ****p < 0.0001.

Gait analysis revealed distinct recovery profiles between adult and young tendons. In injured adult tendons (AT), the injured right paw consistently exhibited a significantly lower angle compared to the contralateral uninjured left paw (Fig. 1C). Although a gradual recovery emerged from week 4, AT never reached the baseline level of uninjured limbs. In contrast, young tendons (YT) showed a rapid recovery as early as week 2, and intriguingly, the right paw angle even surpassed that of the uninjured left paw. This observation suggests a potential growth and adaptation compensation, where developing young rats adopt an altered stepping pattern with an increased outward paw placement during locomotion. Quantitative assessment using the Achilles functional index (AFI ^14^) revealed a similar trend (Fig. 1D). YT exhibited earlier and stronger functional recovery, achieving near-baseline performance by week 4, whereas AT showed a slower trajectory, only partially recovering by week 6.

To evaluate the biomechanical impact of defect healing, contralateral uninjured AT, (injured) AT and YT tendons were subjected to uniaxial tensile testing (Fig. 1E). The cross-sectional area (CSA, the transverse area of the tendon at the mid-substance) of AT increased markedly from 1.52 ± 0.23 mm² to 4.98 ± 0.28 mm², while YT showed a similar increase from 1.42 ± 0.40 mm² to 3.72 ± 0.22 mm². The overall change in the CSA was similar for AT and YT at the endpoint analysis. Failure load, defined by the maximum force a tendon can withstand before rupture, did not fully recover in AT (Fig. 1F). In contrast, YT restored failure load values comparable to uninjured tendons (71.32 **±** 17.37), and normalized comparisons confirmed that YT achieved significantly higher values than AT (p = 0.0008). Despite this higher load-bearing capacity, both AT and YT exhibited significantly reduced tensile strength relative to uninjured tendons, indicating persistent structural weakness in the repaired tendons (Fig. 1G). Within this context, YT displayed significantly greater normalized strength than AT (p = 0.0002). Furthermore, YT (40.39 ± 14.24 MPa) demonstrated a higher degree of elasticity compared to AT (21.65 ± 7.72 MPa), though neither group achieved complete recovery of biomechanical strength or elasticity (Fig. 1H). Taken together, both AT and YT demonstrated stiffer tendon with a wide callus formation compared to uninjured tendon. Moreover, similar load at failure observed for YT with healthy counterpart also suggest that the compensatory thickening occurs at the tendon defect to repair, however, both AT and YT were observed to be biomechanically weaker than uninjured tendons.

Histological analysis at 8 weeks confirmed re-joining of tendon stumps in AT (Fig. 1I). While large areas exhibited re-emerging, longitudinally aligned collagen bundles (green arrows), prominent fibrotic tissue remained (blue arrows), and it was clearly characterized by dense cellular infiltration and disorganized fibers. YT likewise achieved stump bridging (Fig. 1J), but it displayed an expansive fibrosis with neovascularization (red arrow) and disrupted tissue continuity (white arrows). Together with the biomechanical results, these findings indicate that both AT and YT primarily repair via a cellular bridge across the defect followed by deposition of immature ECM. Collectively, these results suggest that young tendons exhibited faster functional recovery and greater mechanical strength than adult tendons at the analyzed time points.

### Major cell populations identified during tendon healing

To delineate the cellular landscape of healing tendon in adult and young animals, we performed single nuclei RNA sequencing (snRNAseq, Fig. 2A). Total readouts of the cell types, cell type-specific gene markers and post-quality contol (post-QC) data are given in supplementary materials (Supp. Tables 1 and 2). Achilles tendon samples were collected at 7, 14-, and 28-days post-injury, corresponding to the acute, proliferative, and remodeling phases of repair and compared to uninjured AT. After merging and integration, dimensional reduction was further corrected using Harmony (v1.2.3), which aligned cells across samples and time points. We identified seven transcriptionally distinct cell populations shared between uninjured tendon and healing tendons (Fig. 2B). The subclusters of the cells were identified according to shared marker genes provided in supplementary materials (Supp. Fig. 1 and 2.)

**Figure 2.**
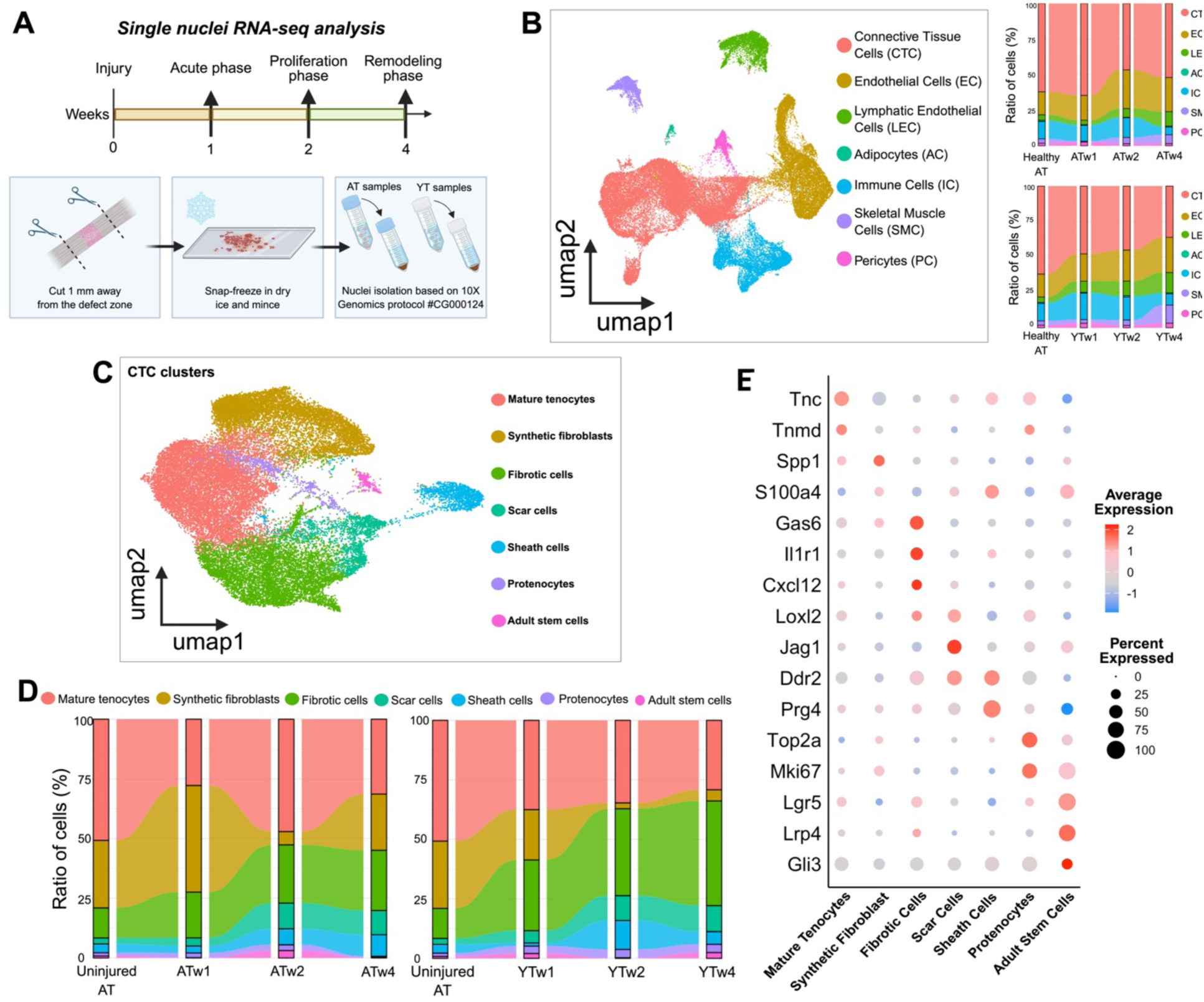
Single nuclei RNA-sequencing revealed the cellular dynamics of major tenogenic and fibroblastic lineages partaking in repair, fibrosis and scar maintenance. **A)** Depiction of single nuclei RNA-sequencing experimental timeline and sample processing across acute, proliferative, and remodeling phases **B)** UMAP of all nuclei resolves major tissue compartments in tendon healing and the quantitative comparison of their numbers throughout the healing. **C)** Sub-setting the connective tissue cells (CTC), seven different cell types were identified as a part of tendon healing program, consisting of tenogenic and fibroblastic lineages. **D)** The dynamic changes within the CTC clusters revealed a striking difference between the adult and young healing modalities, where AT shows an early expansion of synthetic fibroblast population and YT remains biased toward fibrotic and scar-related populations. **E)** Dot plot of marker genes highlights lineage identities and states.

The analysis of cell type composition across uninjured AT, injured AT and YT samples revealed distinct shifts in the relative abundance of major cellular populations during the healing process (Fig. 2B). As the baseline of snRNA-seq data, uninjured AT was used to identify cell types at the homeostatic state. The connective tissue cells (CTCs) dominated the uninjured AT (62.1%), they are identified by a matrix adhesion signature of *Postn^+^/Dcn^+^/Cdh11^+^*(Supp. Fig. 1) typical to tendon stroma^15^. Endothelial cells (EC, 16.2%) were defined by *Flt1* and *Vcam1*^16^. At a much lower quantity, the lymphatic endothelial cells (LEC, 3.8%) were determined by *Prox1^+^/Lyve1^+^/Vegfc*+^17^. Immune cells (IC, 12.2%) expressed *Ptprc* (*CD45*) together with antigen presenting surface receptor *Cd74* ^18^ and T-cell receptor complex *Cd3d.* Pericytes were marked by *Rgs5⁺/Acta2⁺ (αSMA)/Mcam⁺* (*CD146*)^19^. Additional non-CTC (AC) lineages were detected at low levels. Skeletal muscle nuclei showed *Acta1*, *Myh2*, and *Myh7*, reflecting a possible carryover from the myotendinous region^20^.The analysis of cell type composition across uninjured AT, injured AT and YT samples revealed distinct shifts in the relative abundance of major cellular populations during the healing process (Fig. 2B). As the baseline of snRNA-seq data, uninjured AT was used to identify cell types at the homeostatic state. The connective tissue cells (CTCs) dominated the uninjured AT (62.1%), they are identified by a matrix adhesion signature of *Postn^+^/Dcn^+^/Cdh11^+^*(Supp. Fig. 1) typical to tendon stroma^15,21^. Endothelial cells (EC, 16.2%) were defined by *Flt1* ^22^ and *Vcam1*^16^. At a much lower quantity, the lymphatic endothelial cells (LEC, 3.8%) were determined by *Prox1^+^/Lyve1^+^/Vegfc*+. Immune cells (IC, 12.2%) expressed *Ptprc* (*CD45*) together with antigen presenting surface receptor *Cd74* ^18^ and T-cell receptor complex *Cd3d.* Pericytes were marked by *Rgs5⁺/Acta2⁺* (*αSMA)/Mcam⁺* (*CD146*)^19^. Additional non-CTC (AC) lineages were detected at low levels. Skeletal muscle nuclei showed *Acta1*, *Myh2*, and *Myh7*, reflecting a possible carryover from the myotendinous region^20^.

Injured adult tendons demonstrated a marked alteration in their cellular composition over the healing period. At week 1, CTCs remained the largest compartment (64.6%), yet their proportion steadily decreased by week 2 (46.6%) and stabilized around 52.0% at week 4 in AT (Fig. 2B). This decline was accompanied by a relative expansion of ECs and LECs. ECs increased from 17.4% in week 1 to 27.2% in week 2, before slightly declining to 24.0% at week 4. Similarly, LECs expanded progressively, rising from 3.1% to 10.1% over the healing period. ICs, which initially represented 11.1% of the population, displayed a biphasic pattern, increasing modestly at week 2 (13.8%) but then contracting to 5.4% by week 4. AC remained a minor fraction in all adult samples (<1%). Cell populations in the young tendons followed a different trajectory, with notable divergence from the adult response. At week 1, CTCs constituted 47.9% of the sample. This proportion remained stable at week 2 (45.2%) but dropped sharply to 36.2% by week 4. Concomitantly, there was a sustained enrichment of ECs and LECs. In YT, ECs comprised the 19.3% of all populations at week 1, remained consistently elevated at week 2 (21.9%) and further increased to 24.8% at week 4. The most striking expansion occurred within the LEC compartment, which grew from 8.0% at week 1 to 14.4% at week 4. ICs were also more abundant in young tendons during the early phases of healing, reaching 18.8% at week 1 and 16.1% at week 2, before contracting to 8.0% at week 4. Similar to other samples, ACs remained rare (<1%) (Fig. 2B).

### Cellular composition of the adult and young tendon CTC compartment

Sub-setting of the CTC compartment revealed seven transcriptionally distinct clusters that segregated into two major lineages: tenogenic and fibroblastic lineages (Fig. 2C). The technical summary of the nuclei analyzed for CTC subset, highly expressed distinctive genes for each cell type and their distribution among cell clusters are given in supplementary materials (Supp. Fig. 3, supp. table 3). *Tnmd* is a canonical tenocyte marker ^23^ and *Tnc* is a matricellular ECM protein, partaking in tenocyte–matrix adhesion and remodeling^24^. Moreover, the proliferation markers *Mki67* and *Top2a* are typically expressed in actively cycling cells and decline as they exit the cell cycle^25,26^. Mature tenocytes, which accounted for 50.8% of all CTCs in uninjured tendons, were markedly reduced during healing (Fig. 2D). Consistent with this, these cells lacked proliferative markers *Mki67* and *Top2a* but expressed high levels of *Tnmd* and *Tnc,* indicating a quiescent, terminally differentiated state (Fig. 2E). In AT, proportions declined to 27.8% at week 1, partially recovered at 47.1% by week 2, and decreased again to 31.4% at week 4. In contrast, YT exhibited a more progressive loss, falling from 37.5% at week 1 to 23.6% at week 2 and 23.1% at week 4 (Fig. 2D). On the other hand, tenogenic progenitors (protenocytes) were defined as proliferative *Mki67*⁺ / *Top2a*⁺ progenitors with low *Tnmd* and *Tnc*, were rare in uninjured tendon at 1.4% (Fig. 2E). In injured samples, protenocytes rose modestly to 2.0% at week 1 and 2.4% at week 2, before collapsing to 0.5% at week 4 in AT (Fig. 2d). In young tendons, these progenitors were more sustained, expanding to 3.1% at week 1, peaking at 5.9% at week 2, and remaining at 4.9% at week 4.

**Figure 3.**
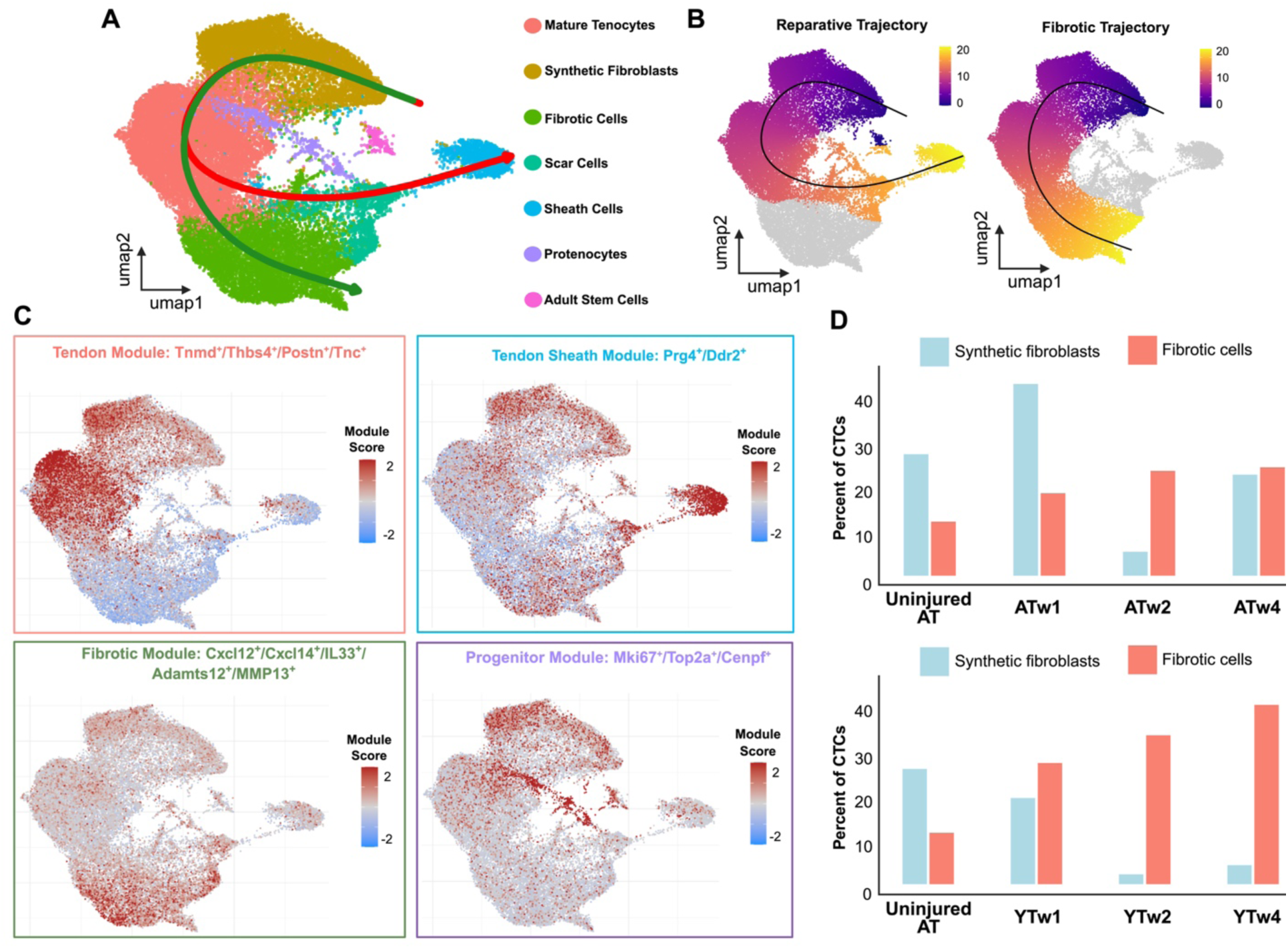
The unsupervised trajectory analysis reveals reparative and fibrotic lineages driving distinct cellular dynamics in tendon healing. **A)** Slingshot trajectory analysis of the CTCs. **B)** Reparative and fibrotic fate branches emerged from the CTCs. **C)** Module scoring UMAPs revealed tendon-sheath and progenitor programs, in addition to two major modules as tenogenic and fibrosis-driving modules. **D)** Quantification of the synthetic fibroblasts and fibrotic cells revealed that adult tendons have preference to maintain regenerative cluster while young tendon tend to expand the fibrotic cluster.

In the fibroblastic lineage, fibrotic cells upregulated *Cxcl12, Il1r1,* and *Mmp2* (Fig. 2E). *Cxcl12* is a strong chemoattractant for leukocytes as well as progenitors^27^. In addition, the *Il1r1* receptor activates downstream *Nk-kB* signaling^28^, and *Mmp2* partakes in matrix breakdown and remodeling^29^. In AT, they have risen from 12.6% to 19.2% at week 1, peaking at 25.2% at week 4 (Fig. 2D). In YT, fibrotic cell expansion was more pronounced, increasing from 29.7% at week 1 to 30.8% at week 2 and reaching 33.9% at week 4.

A distinct ECM-synthetic fibroblast cluster, defined by *S100a4* and *Spp1*, represented 28.3% of CTCs in uninjured tissue (Fig. 2D,E). This cluster is enriched in motility protein *S100a4* ^12^ and in *Spp1*, which takes part in ECM deposition^30^. In AT, these cells surged at week 1 to 44.7% but regressed sharply at week 2 down to 5.6% before stabilizing at 23.5% at week 4. In YT, synthetic fibroblasts decreased over time to 4.7% at week 4, reflecting a striking deactivation of synthetic program.

On the other hand, the MMP-activating fibroblast marker *Ddr2*^31^, collagen crosslinking enzyme *Loxl2* and fibrosis-inducing *Jag1*^32^ characterized the scar-associated fibroblasts (scar cells)(Fig. 2e). In AT, they reached 10.8% at week 2 and 10.1% at week 4 (Fig. 2D). In YT, they followed a similar trajectory but plateaued at slightly higher levels such as 5.2% at week 1, 9.6% at week 2 and 10.5% at week 4.

One of the specialized cell types in tendon are sheath cells were defined by *Prg4* and *Ddr2* (Fig. 2E). *Prg4* takes part in tendon gliding ^33^ and was present in tendon sheath lineage along with *Ddr2*. In AT, this population fluctuated, with 2.9% at week 1, 6.7% at week 2, and 9.0% at week 4 (Fig. 2D). In YT, sheath cells were poorly represented, at 1.3% at week 1 and only 2.6–2.7% at weeks 2 and 4, indicating impaired sheath regeneration. Another specialized cluster, adult stem cells, were identified by their expression of *Lgr5*, *Tgfb1* and *Gli3* (Fig. 2E). *Lgr5* and *Gli3* potentiates *Wnt/β-catenin* signaling, thus sustaining self-renewal^34^. *TGF-β1* directs fate toward matrix-producing phenotypes, such as fibrotic lineages. These cells were rare (0.8%) in uninjured tendon (Fig. 2D).

In AT, they were nearly absent at week 1 (0.1%), expanded transiently to 3.1% at week 2, and declined to 0.2% at week 4. In YT, stem cells were more consistently detected, at 2.0% at week 1, 2.9% at week 2, and 1.0% at week 4.

### Transcriptional trajectories of CTC subsets during tendon healing

To explore the heterogeneity of CTCs during tendon healing, we performed trajectory analyses (Fig. 3A). One of the trajectories showed a reparative path culminating in tendon sheath cells while the other trajectory indicated a fibrotic path (Fig. 3B). The reparative clusters expressed markers which are consistent with matrix production, lubrication, and tendon-sheath interface maintenance. Whereas the pro-fibrotic signals were upregulated in fibrotic lineages (Fig. 3C), such as *Il33*, and *Mmp13* structural damage^35^.

The *Cxcl12* and *Cxcl14* markers demonstrated there is recruitment of fibroblasts that may take part in dysfunctional remodeling^36,37^. Then, the fibrotic program revealed a strong bias towards enrichment of matrix degeneration markers and their co-expression simultaneously (Fig. 3C). We also identified a proliferative, progenitor-like module composed of *Mki67^+^/Top2a^+^/ Cenpf^+^* clusters (Fig. 3C). This revealed a fibroblastic pool that may be poised to supply either reparative sheath cells or fibrotic cells.

Finally, we quantified the relative abundance of synthetic fibroblasts and fibrotic cells across conditions (Fig. 3D). In AT, synthetic fibroblasts predominate early and then contract as fibrotic cells rise by week 4. In YT, fibrotic populations expand earlier and remain dominant across timepoints. This outcome implies that the balance between reparative and fibrotic responses may depend on the shifting proportions of these populations in AT and YT.

### Adult tendon remodeling associated with synthetic fibroblasts

We performed cluster-wise differential expression and visualized marker enrichment to determine the features of the synthetic fibroblast population within the CTC compartment (Fig. 4). An autocrine growth axis via *Pdgfb*^38^ and *Vim, S100a6*, and *Tmsb4x* encoding cellular motility and matrix adhesion^39^ were upregulated in synthetic fibroblasts (Fig. 4A). Furthermore, Spp*1* and *Fabp4* co-expression involving in matrix remodeling and increased fibroblast metabolism^40^ along with *Tpt1* modulator related to survival and proliferation ^41^ were upregulated in these cells.

**Figure 4.**
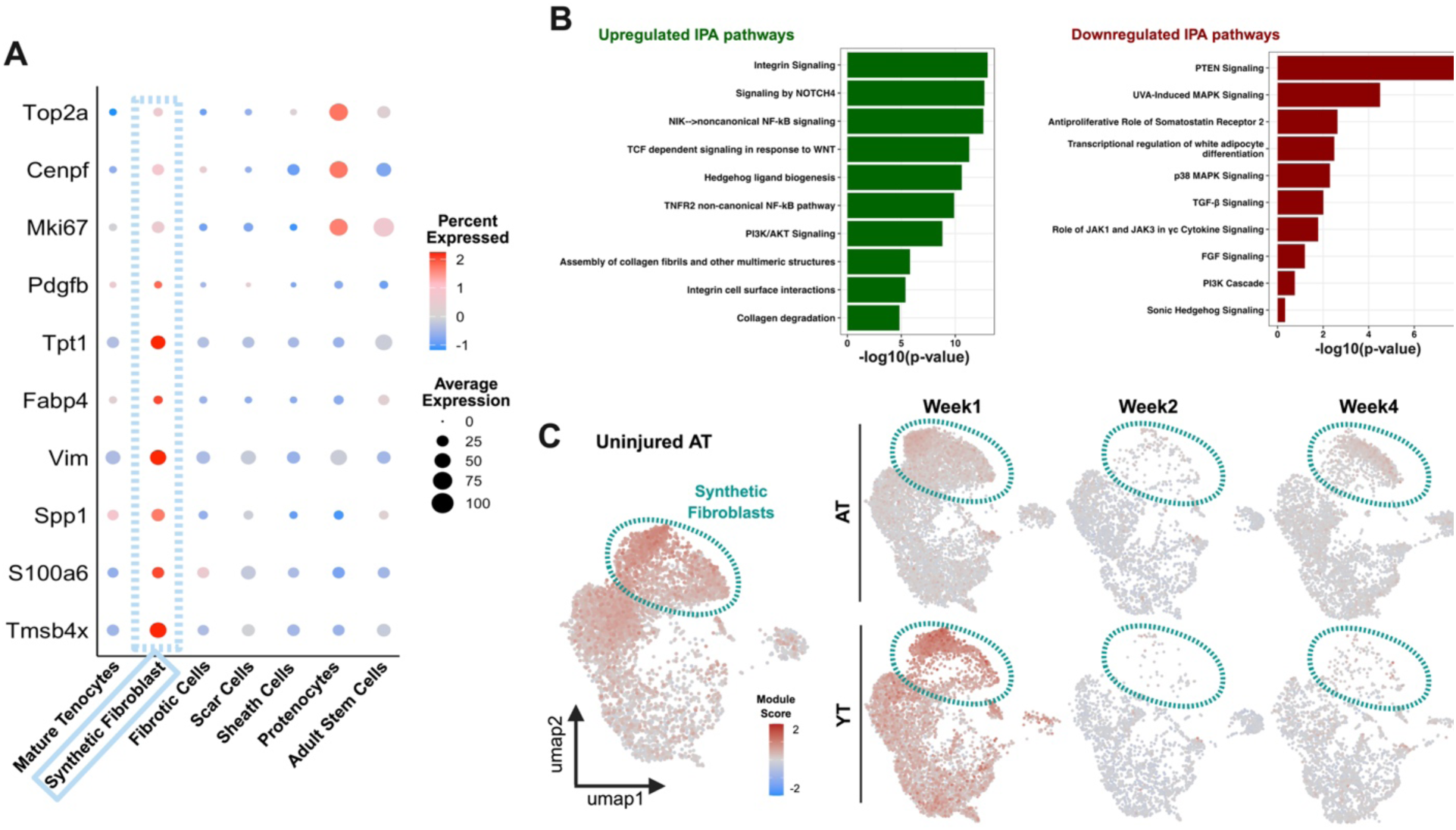
Adult tendons maintained more synthetic fibroblasts in comparison to young tendons. **A)** Synthetic fibroblasts express hallmark genes for ECM remodeling, cell adhesion and maintenance genes. **B)** IPA enrichment indicates upregulation of pathways consistent with reparative remodeling and attenuation of stress/scar-associated signaling. **C)** In AT, the synthetic fibroblast population persists across the healing timeline, whereas in YT it is markedly reduced.

The IPA analysis enriched the genes in relation to Integrin, Wnt/Tcf, Notch, Tnf2/non-canonical Nf-κb, Hedgehog, and collagen assembly terms in synthetic fibroblasts (Fig. 4B). The upregulated pathways were found to converge on cell–matrix adhesion and tissue development^42^. Whereas, the downregulated pathways are dominated by stress-kinase/antiproliferative modules, consistent with a shift toward adhesion-driven growth and matrix production. In AT, the synthetic-fibroblast program surges early (week 1) and maintained through the week 4 (Fig. 4C). In YT, the same program dwindled and decreased significantly over time (Fig. 4D).

### Fibrosis-associated healing in young tendons

To define the fibrotic program within the CTC compartment, we identified key marker genes taking part in fibrotic downstream signaling (Fig. 5A). The gene set consists of chemokine/alarmin, protease, and developmental-receptor genes. The signature combines *Gas6* signaling with pro-fibrotic signaling markers enriched in *TGFβ/activin* receptors such as *Tgfbr2*, *Tgfbr3* and *Acvr2a*^43,44^ and *Wnt* co-receptors *Lrp6* and *Lrp1*. Moreover, the fibrotic cells strongly upregulated *Angpt1* as a vascularization modulator^45^.

**Figure 5.**
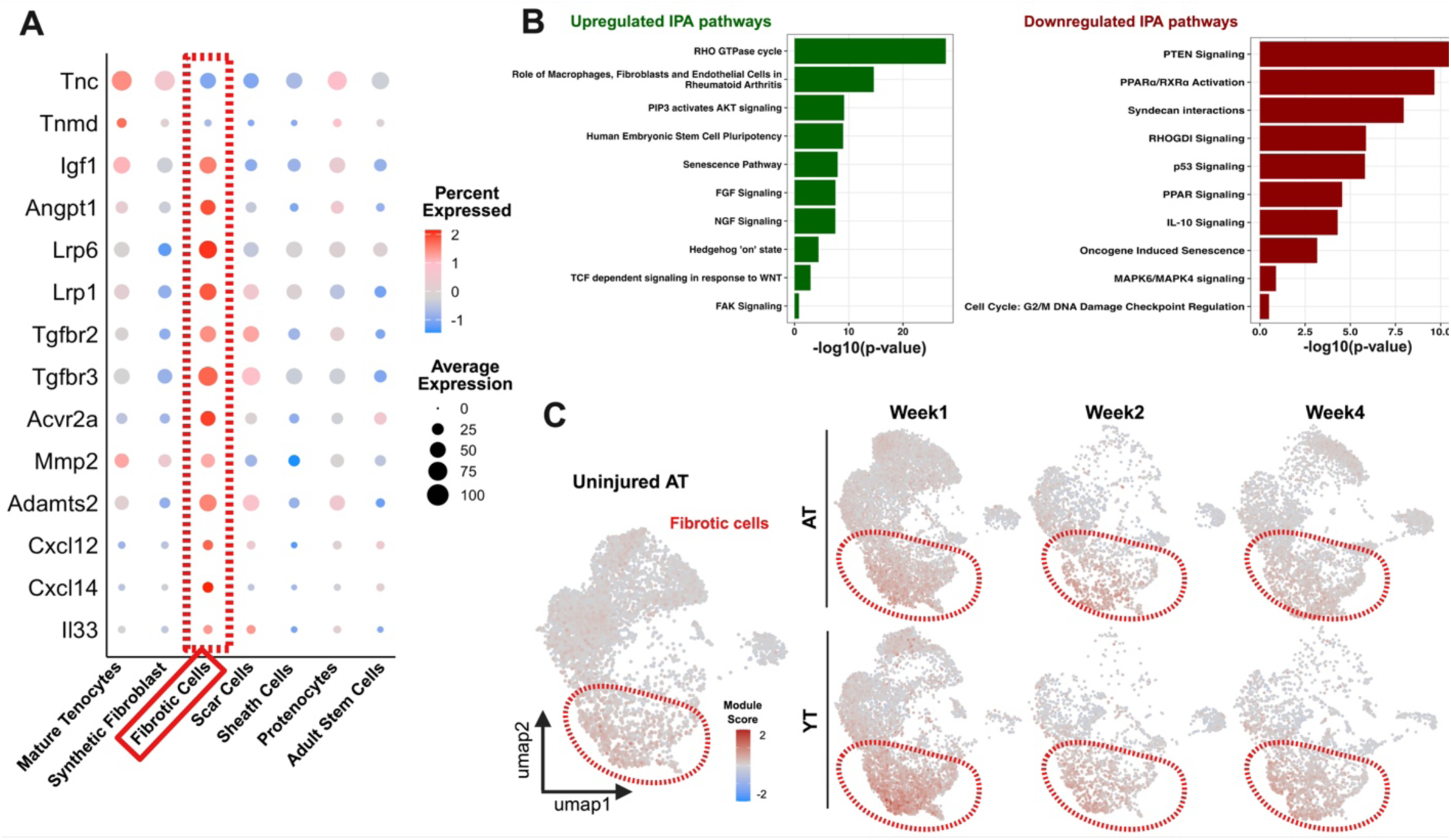
YT responded with higher relative fibrotic and scar cell presence to injury. **A)** Fibrotic cells key genes enriched for pro-fibrotic receptors and co-receptors and ECM-related genes. **B)** IPA analysis shows adhesion and developmental programs leading to fibrosis and downregulation of anti-fibrotic signaling pathways including senescence. **C)** Through the healing timeline, fibrotic cells definitively expanded and persisted in YT.

The IPA analysis (Fig. 5B) showed upregulated response of *Rho GTPase* cycle and *FAK* partaking in increased actomyosin tension and focal-adhesion maturation^46^. Concurrently, *Wnt/Tcf* and *Hedgehog* programs and *TGF-β/activin* receptors were upregulated significantly. Besides, this cluster downregulated *PTEN/PPAR* and stress-checkpoints, such as *p53* and *Il10*, significantly^47^.

Mapping the fibrotic module onto the UMAP (Fig. 5C) and quantifying its compartment size (Fig. 3D) revealed divergent deployment. In AT, the fibrotic program rises transiently during early phases and contracts by later time points. In YT, the same program shows broader and persistent expression of hallmark genes.

### ROS-associated profibrotic signaling in young tendons

*Nox4* and *Gas6* mark a ROS ^48,49^ and survival-related fibrotic signal enrichment that became overactive in fibrotic cells (Fig. 6A). Over time, YT exhibited elevated expression of *Nox4*, accompanied by a marked upregulation of *Vegfa* (Fig. 6B). At the same time, antioxidant gene *Gpx1* expression decreased significantly. In addition, CellChat communication analysis shows amplified bidirectional crosstalk between fibrotic cells and their partners via the information flow from fibrotic cells to overexpressed fibrotic phenotype in YT (Fig. 6C). Furthermore, fibrotic cells acted as highly interactive hubs, with enriched signaling routes involving collagen, laminin, ephrin, VEGF, and THBS pathways (Supp. Fig. 4). As receivers, fibrotic cells were particularly responsive to adhesion and survival-related cues such as *CD44*, *Sdc4*, and *Ngfr*^50,51^. As senders, these cells secreted a broad spectrum of ligands, including *Collagen*, *Laminin*, *Fgf, Angpt,* and *Gas6* (Supp. Fig. 5). Module scoring on UMAPs showed consistently upregulated *Nox4* and *Gas6* signaling in YT at the fibrotic cell-related module (Fig. 6D). In addition, a mechanistic network using IPA centers on ROS production and survival signaling links *Nox4* with *Vegfa, Wnt5a, Lrp1, Map3k5*, and stress-related nodes like *Tlr4, Cxcl12* and *Icam1* and a downregulation in matrix-remodeling outputs such as *Thbs1* and *Spp1* (Fig. 6E).

**Figure 6.**
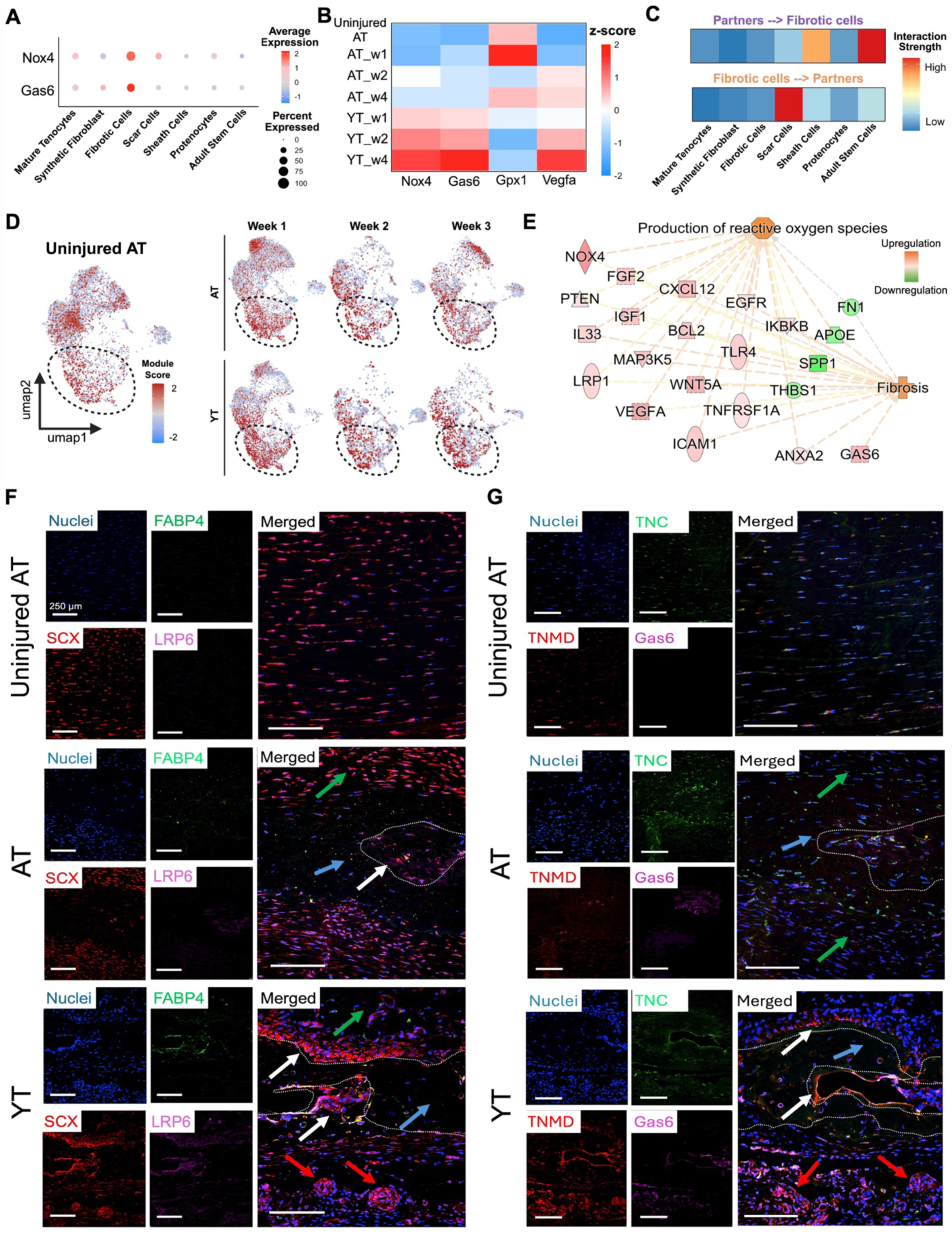
Nox4 and Gas6 may drive fibrosis in tendon healing. **A)** The marker distribution among clusters revealed a dominant Nox4 and Gas6 signaling in fibrotic cells. **B)** Pro-fibrotic Nox4, Gas6 and pro-vasculogenic Vegfa were upregulated in YT over time while Nox4 suppressor Gpx1 decreases in YT. **C)** Fibrotic cells are fed mostly from adult stem cells and sheath cells, and fibrotic cells mostly support scar cells. **D)** Nox4 module scoring localizes enrichment to the fibrotic cluster, consistent with a broad presence of Nox4⁺ cells **E)** A reactive oxygen species (ROS)-centered mechanistic network of differentially regulated genes in fibrotic cells shows a shift toward adhesion/ECM programs and fibrosis. **F)** Uninjured AT has aligned Scx^+^ cells with a very low expression of Fabp4 (synthetic fibroblasts) and Lrp6 (fibrotic cells). AT demonstrated Fabp4 positive cells located at the fibrotic zone (white arrow) and repaired zones (enclosed by white line) devoid of Lrp6 expression (green arrow). Whereas, YT has strong presence of blood vessels (red arrows) with infiltrating Scx^+^ cells and fibrotic scar tissue (white arrow) that is highly Scx^+^/Lrp6^+^. **G)** Uninjured AT shows low Tnmd, sporadic Tnc, and little Gas6 expression. AT follows a similar pattern with a modest rise in Gas6. In contrast, YT reveals well-defined fibrotic niches characterized by Gas6⁺/Tnc⁺ matrix and reduced Tnmd.

Immunofluorescence (IF) imaging revealed an ordered, aligned and highly *Scx*^+^ tenocytes in uninjured tendon (Fig. 6F). *Fabp4* and *Lrp6* expression were essentially absent in homeostatic condition. Injured AT showed *Scx*^+^ alignment in the restored zone (green arrow) around the stabilized scar zone (blue arrow) with low cellularity. A small presence of *Fabp4*^+^ and *Lrp6*^+^ showed reparative synthetic fibroblasts dispersed in the fibrotic area, but it was contained (white arrow). On the contrary, YT had a striking expansion of *Lrp6*^+^ cells throughout the fibrotic area. *Fabp4*^+^ cells were also numerous compared to AT and *Scx*^+^ were fragmented around the defect zone showing that YT has not been able to stabilize the defect zone successfully (white arrows). Moreover, several foci for blood vessel formation have also been detected in YT (red arrows), while these were absent in AT.

To check the presence of maturing tenocytes in both AT and YT, *Tnmd* and *Tnc* proteins were visualized along with *Gas6* (Fig. 6G). Again, uninjured AT demonstrated an aligned structure with minimal *Tnmd* and *Tnc* thus showing an undisturbed and stable tendon structure. However, YT illustrated a highly prominent expression of *Gas6*^+^ and *Tnmd*^+^/*Tnc*^+^ cells around the fibrotic zone.

## Discussion

In this study, we delineate tendon-specific cell populations and signaling programs that govern successful repair versus fibrosis within the connective tissue compartment. Elucidating the cellular mechanisms that distinguish regenerative from fibrotic outcomes is therefore essential for the development of targeted therapies. Accordingly, adult and young tendon groups were selected to represent distinct biological contexts, with YT characterized by a presumed higher abundance of regenerative progenitors, and AT reflecting a more stabilized, homeostatic environment. Using the non-repair injury model across ages allowed us to compare regenerative versus fibrotic transcriptional programs under a consistent injury and natural healing context, while minimizing variability that might be introduced by repair techniques.

Functional analyses showed that young tendons exhibited rapid functional recovery but incomplete structural and molecular restoration, whereas AT tendons healed more slowly yet developed a more organized architecture. Although paw angle measurements and the steeper improvement in the Achilles Functional Index suggested earlier functional improvement in young tendons, this pattern likely reflects the heightened cellularity and active matrix remodeling that accompany postnatal tendon growth, which can accelerate ECM deposition and transiently improve function without representing development-independent regeneration^52^. In addition, a comparable adaptation has been documented in human Achilles tendon healing, where compensatory redistribution of load toward the knee alters ankle motion and angle^53^. An increase in CSA reduces stress for a given load, allowing the tendon to withstand similar forces at lower stress levels^54^. In line with this, the non-repair rat Achilles tendon model here demonstrated that progressive ECM produced a load-bearing yet compositionally altered tissue with reduced elasticity^55^. Similarly, in human anterior cruciate ligament repairs, recovery of load-bearing capacity has been attributed to increased CSA and ECM deposition rather than full restoration of biomechanical features^56^. Consistent with these observations, our findings of comparable load-to-failure but moderate elastic modulus in both AT and YT suggest that mechanical recovery was achieved predominantly through volumetric thickening of the repair tissue, where YT had higher recovery rate than AT.

We analyzed the cell-level differences between injured and uninjured tendon in adult and young tendons through snRNA-seq analysis. The technical metrics and post-QC expression profiles of the samples revealed sufficiently high-quality data for downstream analysis, although there are sequencing depth variances. In addition, among the annotated populations of cells, connective tissue cells showed particularly robust transcriptomic support. CTC was represented by a large number of nuclei and displayed strong expression of canonical extracellular matrix genes, including Col1a1, Col3a1, and Fn1, each detected across a high proportion of nuclei.

Interestingly, YT recovered biomechanical features faster and more prominently than AT. However, we found that the adult group exhibited a more restrained remodeling response, characterized by expansion of synthetic fibroblasts but a limited and well-contained fibrotic plaque. Our data illustrated that YT revealed a less regulated healing which was characterized by disorganized remodeling and incomplete structural restoration. In addition, YT displayed a persistent expansion of fibrotic populations accompanied by progressive erosion of the tenogenic niche. Although *Lgr5⁺/Lrp4⁺/Gli3⁺* adult stem-like cells persisted as a small pool in both age groups, their prevalence was markedly higher in YT, suggesting a greater propensity for lineage diversion. In agreement with prior studies identifying *Lgr5⁺* mesenchymal progenitors as contributors to musculoskeletal tissue formation^57^, our data indicate that, under dysregulated signaling conditions, these cells may undergo fibrotic reprogramming, as evidenced by co-expression of *Lgr5* and *Lrp4*. The enrichment of *Lrp6⁺*or *Gas6⁺* cells within the fibrotic cluster further supports activation of a canonical *Wnt/β-catenin–TCF* axis driving maladaptive connective tissue remodeling^58,59^. As this mechanism has not been previously reported in tendon fibrosis, we propose that sustained *Wnt-*dependent *Lrp6* activation may restrict the differentiation capacity of adult stem cells, creating a feed-forward fibrotic loop that preserves the fibrotic cluster and hinders regenerative tendon remodeling. This phenomenon aligns with emerging concepts of a “pro-fibrotic lineage” enriched in *Gas6⁺/Lrp6⁺* signaling hubs and sustained by the *Lgr5⁺/Lrp4⁺/Gli3⁺* niche, collectively creating a self-stabilizing, pro-adhesive microenvironment that impedes regenerative remodeling. Pathway enrichment further revealed suppression of *Pten, Ppar,* and *Il-10*, alongside reduced p53 signaling, consistent with a transition toward a self-reinforcing fibrotic state^60^. Collectively, these findings delineate a *Wnt-Lrp6*–driven fibrotic program in which adult stem-like populations may be trapped in a fibrotic feed-forward circuit.

In contrast, AT displayed a transient activation of progenitors and *Fabp4⁺/S100a6⁺/Vim⁺* synthetic fibroblasts early in healing. This subset represented a reparative fibroblastic lineage actively remodeling the extracellular matrix, with low but detectable *Mki67⁺/Top2a⁺/Cenpf⁺* expression indicative of controlled cycling and non-fibrotic behavior. Dividing and migrating cell populations such as *Scx⁺* tenogenic cells and epitenon-derived progenitors ^61^ have similarly been shown to contribute to tendon regeneration, though their proliferative potential appears limited. In parallel, *Prg4⁺*sheath fibroblasts maintained a lubricative, reparative phenotype yet underwent limited proliferative expansion, implying that while the gliding interface was preserved in AT more prominently than YT. The persistence of these *Prg4⁺* populations in a low-proliferation, low-inflammatory state underscores the restricted regenerative drive during natural tendon repair.

These observations suggest that YT may undergo a form of regenerative fibrosis, in which rapid fibrotic matrix deposition initially supports tissue continuity and biomechanical recovery without achieving true structural regeneration. In this context, fibrosis may provide an early functional advantage by stabilizing the injured tendon, yet remain qualitatively distinct from native tendon restoration because it is accompanied by disorganized remodeling, persistent fibrotic cell expansion, and progressive loss of the tenogenic niche. Thus, YT appears to occupy an intermediate healing state in which fibrosis is not purely maladaptive, but only partially regenerative and ultimately incomplete.

Mechanistically, fibrotic populations displayed a pronounced upregulation of the *Nox4–Gas6* module, accompanied by increased *Vegfa* and *Angpt1* expression, defining an oxidative, angiogenic, and highly reactive fibroblast phenotype^62^. The coexistence of *Lrp6⁺/Gas6⁺* zones within vascularized, disorganized fibrotic tissue supports the view that *Nox4* acts as a key effector of fibrosis, consistent with its described role in multiple soft-tissue fibrotic contexts^63^. In this work, nox4 and Gas6 were prioritized because they captured a fibrotic program that may initially support rapid tissue bridging and early biomechanical recovery. In this context, the Nox4–Gas6 axis may reflect a form of regenerative fibrosis, in which oxidative, pro-survival, and pro-angiogenic signaling temporarily facilitates functional restoration after injury. Immunofluorescence corroborated these findings, showing well-aligned *Scx⁺* cells in zones with minimal *Fabp4 and Lrp6* expression in AT, whereas YT exhibited prominent *Gas6⁺/Lrp6⁺* fibrotic foci interspersed with vascular ingrowth. Together, these data position *Nox4⁺/Gas6⁺* fibroblasts at the intersection of oxidative stress, angiogenesis, and maladaptive ECM accumulation.

The study highlights a pivotal relationship between fibroblast lineage and the balance between reparative and fibrotic outcomes in tendon healing, linking oxidative stress dynamics to cell-fate specification during tendon repair. A limitation of the study is the partial clinical relevance of the non-repair model. In humans, full tears are usually surgically repaired. However, we have decided not introduce variability related to the surgical technique and explore the natural repair process and its cellular mechanisms. Another limitation of this study is that the young cohort represents a postnatal developmental stage; therefore, age-associated differences in the injury response may reflect developmental state in addition to healing mechanisms. Because young rats are still developing, the data has been interpreted as context-dependent cell-state programs revealed by a biological contrast, rather than as a development-independent measure of intrinsic healing capacity to identify connective tissue cells partaking in tendon healing. While our findings reveal a potential mechanistic link between redox signaling and fibrotic remodeling, the study is limited by the absence of direct *Nox4* or *Gas6* functional validation, and the snapshot nature of the temporal sampling. Therefore, a direct inhibition or genetic manipulation of *Nox4* approach and observation through temporal changes of the lineages would be necessary to establish causality.

## Methods

### Tendon injury

For all studies, female young (3 weeks) and adult (18-20 weeks) Sprague Dawley rats (SD 400 Charles River) were anesthetized under isoflurane (4%) on thermal pads. After shaving the right hind leg, skin surface was sterilized using 1% hydrogen peroxide in PBS and betadine and 1 cm skin incision was made. The Achilles tendon was isolated from surrounding tissue using curved forceps and a full thickness tendon resection without repair was performed at the midsubstance using no. 15 blade. Further details in supplemental information.

### Gait Analysis

Non-toxic red and blue inks were applied to the fore and hind paws of rats, and each animal was placed at one end of a white paper-lined corridor (approximately 10 cm × 48 cm). Prints were collected before surgery as baseline and at 3 days, and 2, 4, 6 and 8 weeks post-injury (n=11). The paw prints were scanned and analyzed using Fiji ImageJ to measure stride length, stride width, paw angle, heel length, and paw width. Further details in supplemental information.

### Biomechanical Analysis

Adult and young tendons were harvested after 8 weeks of healing (n=9). Following dissection from muscle and calcaneus, Achilles tendons were carefully cleaned of surrounding muscle tissue. The central segment approximately 3 mm above and below the defect site was isolated for analysis. The diameters of the isolated tendons were measured with a help of caliper. Further details in supplemental information.

### Histology and immunofluorescence

After gait analysis, randomly selected rats were euthanized (n=3). The right hind leg Achilles tendons from young and adult rats were harvested as samples and the contralateral uninjured Achilles tendons served as controls. Further details in supplemental information.

### Single Nuclei RNA Sequencing

The tendons were carefully harvested and the midsubtances were isolated. For each group, 6 tendons were pooled, placed in 1.5 mL centrifuge tubes in the dry ice and snap-frozen. Then, tendons were moved to wet ice, thermally equilibrated for 5 min and the nuclei isolation buffer prepared using the 10X genomics protocol #CG000124 were added up to 1 mL. Tendons were minced with surgical scissors for 15 s and digested for 3 min. Following digestion, nuclei and debris suspension were double filtered through 30 mm filters (Miltenyi Biotec) and washed once using phosphate buffered saline (PBS, 0.01 M, pH 7.4). Further details in supplemental information.

### Statistical analysis

All statistical analyses, except for SnRNAseq, were performed via Prism 10 (GraphPad). Parametric analyses such as biomechanical tests were analyzed with one-way ANOVA. For multiple comparisons^64^, appropriate post hoc tests were used. Further details in supplemental information.

## Acknowledgements

This work was supported in part by the Cedars-Sinai Applied Genomics, Computational and Translational Core, Cedars-Siani Biobehavioral Core and the Cedars-Sinai Biobank and Research Pathology Resource.

## Author Contributions

A.E.P, M.B. and J.S. conducted the experiments; A.E.P, M.B., M.O., O.S., D.R., L.Z., M.M., W.T., D.H. analyzed the results; A.E.P, M.B., O.S. and D.S. wrote the manuscript.

## Resource availability

### Lead contact

Further information and requests for resources should be directed to and will be fulfilled by the lead contact, Dr. Dmitriy Sheyn.

## Materials availability

This study did not generate new unique reagents.

## Data availability

Any information required about the data reported in this paper is available from the lead contact upon request.

