## Supplemental Information for "Age-Dependent Fibroblast Programs Govern Regenerative and Fibrotic Tendon Repair"

**Supplementary Table 1.** Pre-QC technical summary across samples.

| Sample | Estimated number of cells | Mean reads per cell | Median UMI counts per cell | Sequencing saturation |
|---|---|---|---|---|
| Uninjured | 13978 | 103254 | 1363 | 77.6% |
| Adult week 1 | 13340 | 91873 | 1264 | 63.6% |
| Adult week 2 | 5497 | 269361 | 1856 | 76.7% |
| Adult week 4 | 8118 | 140711 | 964 | 86.1% |
| Young week 1 | 14824 | 70064 | 2776 | 68.0% |
| Young week 2 | 4335 | 308973 | 1695 | 79.2% |
| Young week 4 | 6692 | 197885 | 1493 | 88.9% |

**Supplementary Table 2.** Technical summary of the cell type readouts post-QC.

**Adipocytes**

| Rank | Gene | CPM | Pseudobulk UMI | Avg UMI / nucleus | % nuclei expressing |
|---|---|---|---|---|---|
| 1 | AY172581.24 | 30,051.06 | 19,888 | 54.79 | 99.2% |
| 2 | Ghr | 14,279.09 | 9,450 | 26.03 | 95.9% |
| 3 | Ebf1 | 8,739.71 | 5,784 | 15.93 | 99.2% |
| 4 | AY172581.9 | 6,299.42 | 4,169 | 11.48 | 86.0% |
| 5 | Kcnip1 | 6,145.30 | 4,067 | 11.20 | 84.3% |
| 6 | Col1a1 | 4,721.92 | 3,125 | 8.61 | 97.2% |
| 7 | Nrxn1 | 4,585.93 | 3,035 | 8.36 | 87.6% |
| 8 | Pdzrn3 | 4,525.49 | 2,995 | 8.25 | 92.6% |
| 9 | Zbtb20 | 3,973.97 | 2,630 | 7.25 | 94.8% |
| 10 | Nfia | 3,213.93 | 2,127 | 5.86 | 88.4% |

**Connective Tissue Cells**

| Rank | Gene | CPM | Pseudobulk UMI | Avg UMI / nucleus | % nuclei expressing |
|---|---|---|---|---|---|
| 1 | AY172581.24 | 39,248.92 | 2,694,429 | 84.29 | 99.8% |
| 2 | AY172581.9 | 8,777.66 | 602,584 | 18.85 | 91.9% |
| 3 | Col1a1 | 8,536.42 | 586,023 | 18.33 | 98.6% |
| 4 | Col3a1 | 6,596.88 | 452,874 | 14.17 | 96.6% |
| 5 | Ebf1 | 6,017.75 | 413,117 | 12.92 | 97.3% |
| 6 | Mt-co2 | 4,804.11 | 329,801 | 10.32 | 78.2% |
| 7 | Mt-co1 | 4,077.49 | 279,919 | 8.76 | 77.9% |
| 8 | Mt-cox3 | 3,895.37 | 267,416 | 8.37 | 80.0% |
| 9 | Fn1 | 3,442.14 | 236,302 | 7.39 | 90.8% |
| 10 | Mt-cytb | 3,292.29 | 226,015 | 7.07 | 69.3% |

**Endothelial Cells**

| Rank | Gene | CPM | Pseudobulk UMI | Avg UMI / nucleus | % nuclei expressing |
|---|---|---|---|---|---|
| 1 | AY172581.24 | 38,426.24 | 1,499,738 | 120.97 | 99.7% |
| 2 | AY172581.9 | 8,536.52 | 333,172 | 26.87 | 89.8% |
| 3 | Mt-co2 | 7,123.96 | 278,041 | 22.43 | 78.5% |
| 4 | Mt-co1 | 5,825.67 | 227,370 | 18.34 | 78.3% |
| 5 | Ebf1 | 5,535.06 | 216,028 | 17.42 | 98.6% |
| 6 | Mt-cox3 | 5,198.62 | 202,897 | 16.37 | 79.7% |
| 7 | Mt-atp6 | 4,437.37 | 173,186 | 13.97 | 77.5% |
| 8 | Mt-cytb | 4,378.82 | 170,901 | 13.78 | 72.4% |
| 9 | Tcf4 | 4,165.70 | 162,583 | 13.11 | 96.7% |
| 10 | Cdh13 | 3,186.68 | 124,373 | 10.03 | 90.5% |

**Immune Cells**

| Rank | Gene | CPM | Pseudobulk UMI | Avg UMI / nucleus | % nuclei expressing |
|---|---|---|---|---|---|
| 1 | AY172581.24 | 54,805.32 | 750,220 | 100.27 | 99.9% |
| 2 | AY172581.9 | 12,498.38 | 171,088 | 22.87 | 93.8% |
| 3 | Mt-co2 | 6,723.15 | 92,032 | 12.30 | 86.8% |
| 4 | Col1a1 | 6,377.62 | 87,302 | 11.67 | 96.3% |
| 5 | Mt-co1 | 6,272.71 | 85,866 | 11.48 | 87.1% |
| 6 | Mt-cox3 | 5,480.09 | 75,016 | 10.03 | 86.7% |
| 7 | Zeb2 | 4,231.48 | 57,924 | 7.74 | 90.2% |
| 8 | Mt-atp6 | 4,215.12 | 57,700 | 7.71 | 83.3% |
| 9 | Mt-cytb | 3,677.09 | 50,335 | 6.73 | 78.7% |
| 10 | Col3a1 | 3,408.04 | 46,652 | 6.24 | 90.3% |

**Lymphatic Endothelial Cells**

| Rank | Gene | CPM | Pseudobulk UMI | Avg UMI / nucleus | % nuclei expressing |
|---|---|---|---|---|---|
| 1 | AY172581.24 | 34,041.37 | 458,045 | 104.72 | 99.5% |
| 2 | Ccl21 | 10,834.50 | 145,784 | 33.33 | 53.6% |
| 3 | AY172581.9 | 7,949.51 | 106,965 | 24.45 | 85.2% |
| 4 | Mt-co2 | 5,952.12 | 80,089 | 18.31 | 70.8% |
| 5 | Mt-co1 | 4,903.93 | 65,985 | 15.09 | 70.9% |
| 6 | Mt-cox3 | 4,218.26 | 56,759 | 12.98 | 72.0% |
| 7 | Mt-cytb | 4,186.38 | 56,330 | 12.88 | 62.5% |
| 8 | Pde7b | 4,178.13 | 56,219 | 12.85 | 91.4% |
| 9 | Mt-atp6 | 3,767.67 | 50,696 | 11.59 | 68.6% |
| 10 | Col1a1 | 3,057.63 | 41,142 | 9.41 | 93.5% |

**Pericytes**

| Rank | Gene | CPM | Pseudobulk UMI | Avg UMI / nucleus | % nuclei expressing |
|---|---|---|---|---|---|
| 1 | AY172581.24 | 50,352.26 | 186,825 | 130.56 | 99.4% |
| 2 | AY172581.9 | 12,533.02 | 46,502 | 32.50 | 90.1% |
| 3 | Ebf1 | 7,424.08 | 27,546 | 19.25 | 99.3% |
| 4 | Mt-co2 | 7,148.90 | 26,525 | 18.54 | 81.3% |
| 5 | Mt-co1 | 6,713.90 | 24,911 | 17.41 | 81.3% |
| 6 | Col1a1 | 6,654.07 | 24,689 | 17.25 | 96.6% |
| 7 | Mt-atp6 | 5,275.50 | 19,574 | 13.68 | 81.3% |
| 8 | Mt-cox3 | 5,265.53 | 19,537 | 13.65 | 82.2% |
| 9 | Mt-cytb | 5,179.55 | 19,218 | 13.43 | 74.9% |
| 10 | Pde3a | 4,337.31 | 16,093 | 11.25 | 93.4% |

**Skeletal Muscle Cells**

| Rank | Gene | CPM | Pseudobulk UMI | Avg UMI / nucleus | % nuclei expressing |
|---|---|---|---|---|---|
| 1 | Neb | 21,940.26 | 120,587 | 48.92 | 98.8% |
| 2 | Dmd | 19,847.52 | 109,085 | 44.25 | 97.2% |
| 3 | AY172581.24 | 12,311.34 | 67,665 | 27.45 | 99.4% |
| 4 | Trdn | 10,050.12 | 55,237 | 22.41 | 96.4% |
| 5 | Mybpc1 | 5,594.46 | 30,748 | 12.47 | 83.4% |
| 6 | AABR07052585.1 | 5,579.18 | 30,664 | 12.44 | 90.9% |
| 7 | Zbtb20 | 5,043.53 | 27,720 | 11.25 | 95.8% |
| 8 | Rbfox1 | 4,794.26 | 26,350 | 10.69 | 90.2% |
| 9 | Mef2c | 3,834.68 | 21,076 | 8.55 | 90.7% |
| 10 | Cdh13 | 3,496.26 | 19,216 | 7.80 | 92.3% |

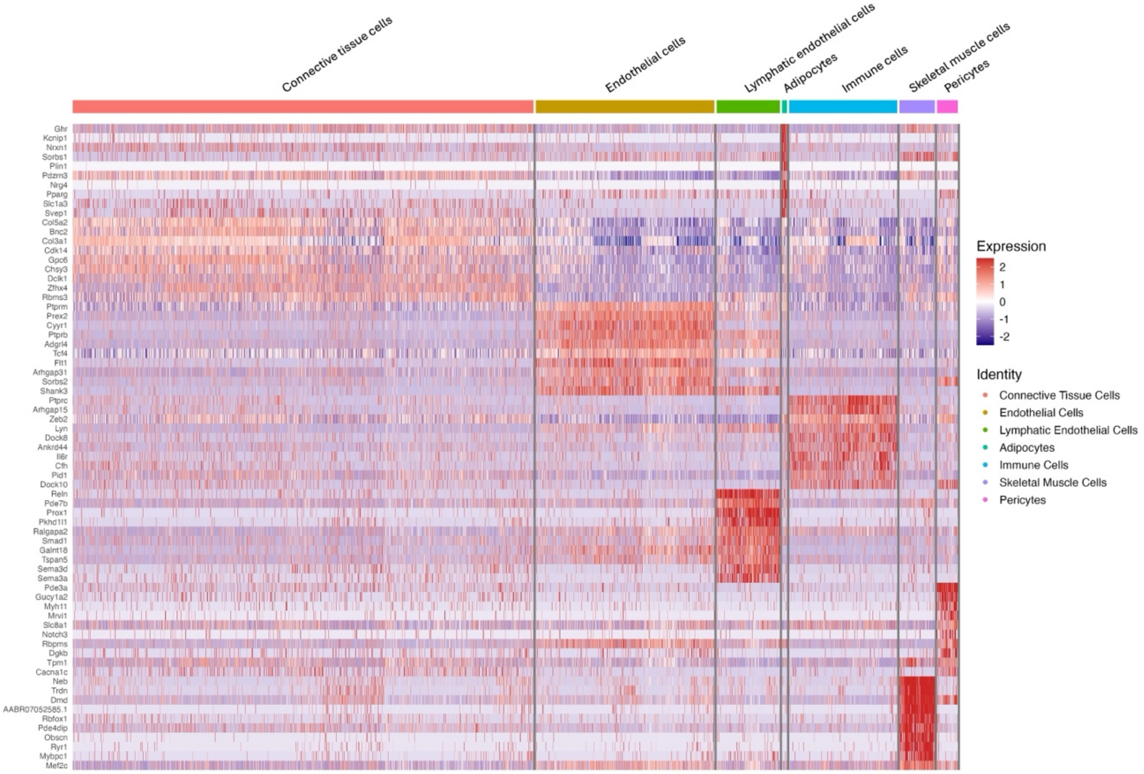


**Supplementary Figure S1.** Heatmap of distinct marker genes for the cell types partaking in tendon healing.

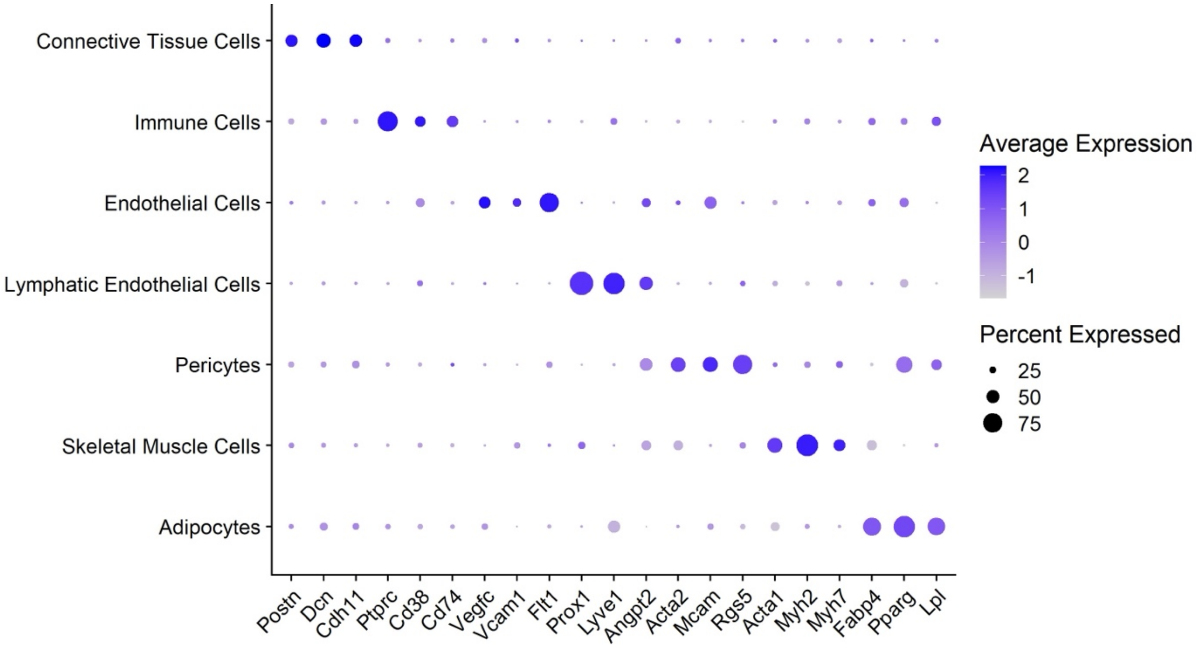


**Supplementary Figure S2.** Key marker genes based on expression in distinct cell compartments in tendon healing.

**Supplementary Table 3.** Technical summary of nuclei analyzed in CTC compartment.

**Mature Tenocytes**

| Rank | Gene | CPM | Pseudobulk UMI | Avg UMI / nucleus | % nuclei expressing |
|---|---|---|---|---|---|
| 1 | AY172581.24 | 32,147.62 | 788,860 | 68.18 | 99.8% |
| 2 | Col1a1 | 8,222.25 | 201,763 | 17.44 | 99.2% |
| 3 | AY172581.9 | 6,868.10 | 168,534 | 14.57 | 91.8% |
| 4 | Col3a1 | 5,677.73 | 139,324 | 12.04 | 96.5% |
| 5 | Ebf1 | 5,455.27 | 133,865 | 11.57 | 97.9% |
| 6 | Postn | 4,690.31 | 115,094 | 9.95 | 89.4% |
| 7 | Col5a2 | 3,749.22 | 92,001 | 7.95 | 94.8% |
| 8 | Trps1 | 3,642.01 | 89,370 | 7.72 | 94.0% |
| 9 | Tnc | 3,595.18 | 88,221 | 7.62 | 84.6% |
| 10 | Fn1 | 3,439.18 | 84,393 | 7.29 | 91.8% |

**Synthetic Fibroblasts**

| Rank | Gene | CPM | Pseudobulk UMI | Avg UMI / nucleus | % nuclei expressing |
|---|---|---|---|---|---|
| 1 | AY172581.24 | 78,151.51 | 775,850 | 96.13 | 100.0% |
| 2 | Hba-a3 | 20,233.57 | 200,869 | 24.89 | 12.4% |
| 3 | LOC100134871 | 17,995.75 | 178,653 | 22.14 | 10.9% |
| 4 | AY172581.9 | 17,404.56 | 172,784 | 21.41 | 96.9% |
| 5 | Col1a1 | 12,533.86 | 124,430 | 15.42 | 99.4% |
| 6 | LOC103694857 | 9,379.29 | 93,113 | 11.54 | 9.6% |
| 7 | Hba-a2 | 8,096.19 | 80,375 | 9.96 | 9.6% |
| 8 | Col3a1 | 6,939.10 | 68,888 | 8.54 | 97.7% |
| 9 | Ebf1 | 3,797.93 | 37,704 | 4.67 | 95.5% |
| 10 | AABR07000398.1 | 3,611.48 | 35,853 | 4.44 | 93.6% |

**Fibrotic Cells**

| Rank | Gene | CPM | Pseudobulk UMI | Avg UMI / nucleus | % nuclei expressing |
|---|---|---|---|---|---|
| 1 | AY172581.24 | 34,854.75 | 685,865 | 90.54 | 99.6% |
| 2 | Col1a1 | 8,750.16 | 172,184 | 22.73 | 97.9% |
| 3 | AY172581.9 | 8,101.72 | 159,424 | 21.05 | 89.1% |
| 4 | Col3a1 | 7,829.84 | 154,074 | 20.34 | 96.7% |
| 5 | Ebf1 | 5,512.50 | 108,474 | 14.32 | 98.5% |
| 6 | Zbtb20 | 3,180.99 | 62,595 | 8.26 | 96.7% |
| 7 | Plxdc2 | 2,867.60 | 56,428 | 7.45 | 91.2% |
| 8 | Col5a2 | 2,842.69 | 55,938 | 7.38 | 92.1% |
| 9 | Fbn1 | 2,758.54 | 54,282 | 7.17 | 87.2% |
| 10 | Rbms3 | 2,755.08 | 54,214 | 7.16 | 93.8% |

**Scar Cells**

| Rank | Gene | CPM | Pseudobulk UMI | Avg UMI / nucleus | % nuclei expressing |
|---|---|---|---|---|---|
| 1 | AY172581.24 | 28,983.49 | 162,937 | 85.62 | 99.6% |
| 2 | Ebf1 | 11,667.96 | 65,594 | 34.47 | 99.7% |
| 3 | Col3a1 | 9,033.18 | 50,782 | 26.69 | 97.7% |
| 4 | Fbn1 | 8,359.37 | 46,994 | 24.69 | 95.9% |
| 5 | Col1a1 | 6,868.90 | 38,615 | 20.29 | 97.7% |
| 6 | AY172581.9 | 6,664.87 | 37,468 | 19.69 | 86.7% |
| 7 | Fn1 | 5,348.90 | 30,070 | 15.80 | 93.9% |
| 8 | Ext1 | 4,156.56 | 23,367 | 12.28 | 96.7% |
| 9 | Cdh13 | 3,437.38 | 19,324 | 10.15 | 87.7% |
| 10 | Creb5 | 3,121.29 | 17,547 | 9.22 | 92.4% |

**Sheath Cells**

| Rank | Gene | CPM | Pseudobulk UMI | Avg UMI / nucleus | % nuclei expressing |
|---|---|---|---|---|---|
| 1 | AY172581.24 | 40,337.77 | 149,761 | 102.79 | 99.8% |
| 2 | Ebf1 | 12,331.54 | 45,783 | 31.42 | 99.5% |
| 3 | Fn1 | 10,358.30 | 38,457 | 26.39 | 97.7% |
| 4 | AY172581.9 | 9,462.72 | 35,132 | 24.11 | 85.7% |
| 5 | Creb5 | 4,547.40 | 16,883 | 11.59 | 91.2% |
| 6 | Sox5 | 3,883.72 | 14,419 | 9.90 | 89.4% |
| 7 | Trps1 | 3,685.48 | 13,683 | 9.39 | 95.4% |
| 8 | Cdh13 | 3,381.66 | 12,555 | 8.62 | 88.3% |
| 9 | Sema5a | 3,366.85 | 12,500 | 8.58 | 83.1% |
| 10 | Zeb2 | 3,321.06 | 12,330 | 8.46 | 91.4% |

**Protenocytes**

| Rank | Gene | CPM | Pseudobulk UMI | Avg UMI / nucleus | % nuclei expressing |
|---|---|---|---|---|---|
| 1 | AY172581.24 | 32,913.00 | 74,917 | 111.82 | 99.9% |
| 2 | AY172581.9 | 7,823.96 | 17,809 | 26.58 | 93.1% |
| 3 | Col1a1 | 7,018.24 | 15,975 | 23.84 | 98.7% |
| 4 | Ebf1 | 6,820.54 | 15,525 | 23.17 | 98.7% |
| 5 | Col3a1 | 6,473.91 | 14,736 | 21.99 | 99.1% |
| 6 | Col5a2 | 3,858.60 | 8,783 | 13.11 | 97.3% |
| 7 | Trps1 | 3,348.10 | 7,621 | 11.37 | 95.4% |
| 8 | Fn1 | 3,306.37 | 7,526 | 11.23 | 93.9% |
| 9 | Cdk14 | 2,849.47 | 6,486 | 9.68 | 95.8% |
| 10 | Gpc6 | 2,624.10 | 5,973 | 8.91 | 88.1% |

**Adult Stem Cells**

| Rank | Gene | CPM | Pseudobulk UMI | Avg UMI / nucleus | % nuclei expressing |
|---|---|---|---|---|---|
| 1 | AY172581.24 | 32,892.94 | 21,640 | 65.38 | 99.4% |
| 2 | AY172581.9 | 6,887.15 | 4,531 | 13.69 | 86.4% |
| 3 | Col1a1 | 5,955.38 | 3,918 | 11.84 | 96.4% |
| 4 | Tchh | 4,439.94 | 2,921 | 8.82 | 18.7% |
| 5 | Pard3 | 3,918.58 | 2,578 | 7.79 | 74.0% |
| 6 | Arl15 | 3,655.62 | 2,405 | 7.27 | 79.2% |
| 7 | Col3a1 | 3,287.77 | 2,163 | 6.53 | 86.7% |
| 8 | Pdzrn3 | 2,568.81 | 1,690 | 5.11 | 67.7% |
| 9 | Cux1 | 2,457.85 | 1,617 | 4.89 | 66.8% |
| 10 | Bnc2 | 2,333.21 | 1,535 | 4.64 | 75.2% |

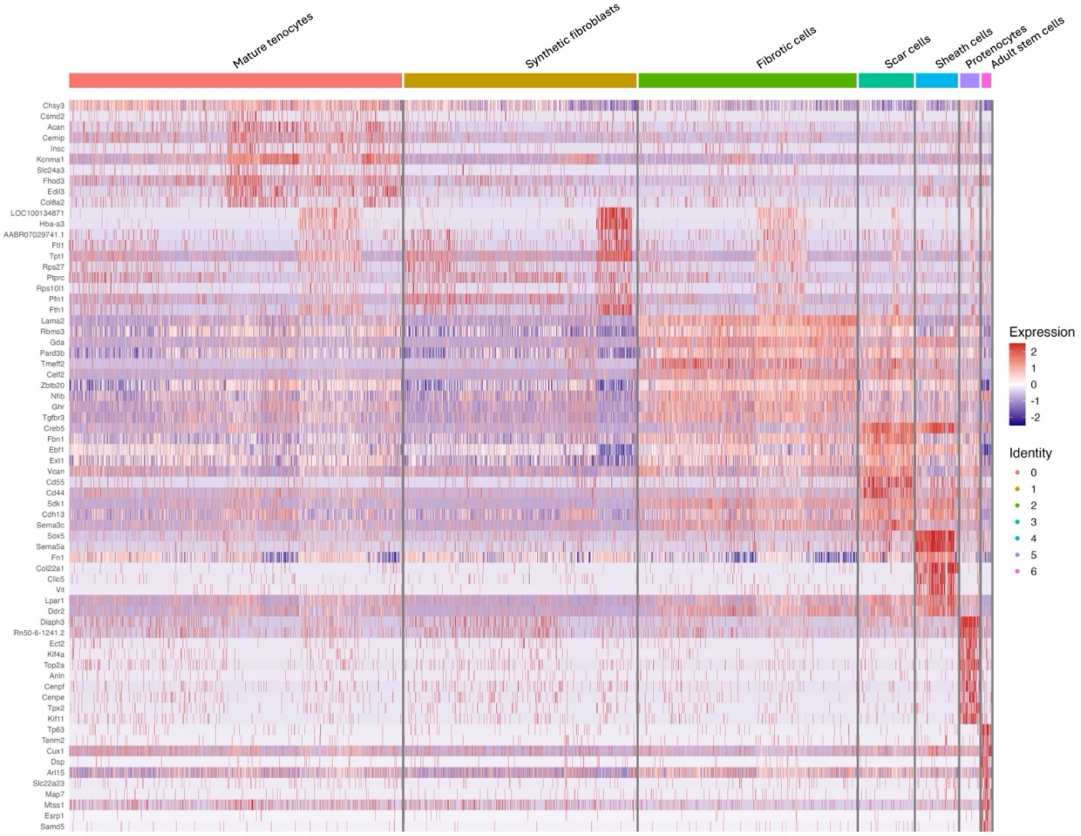


**Supplementary Figure S3.** Heatmap of marker genes specific to each connective tissue cell cluster.

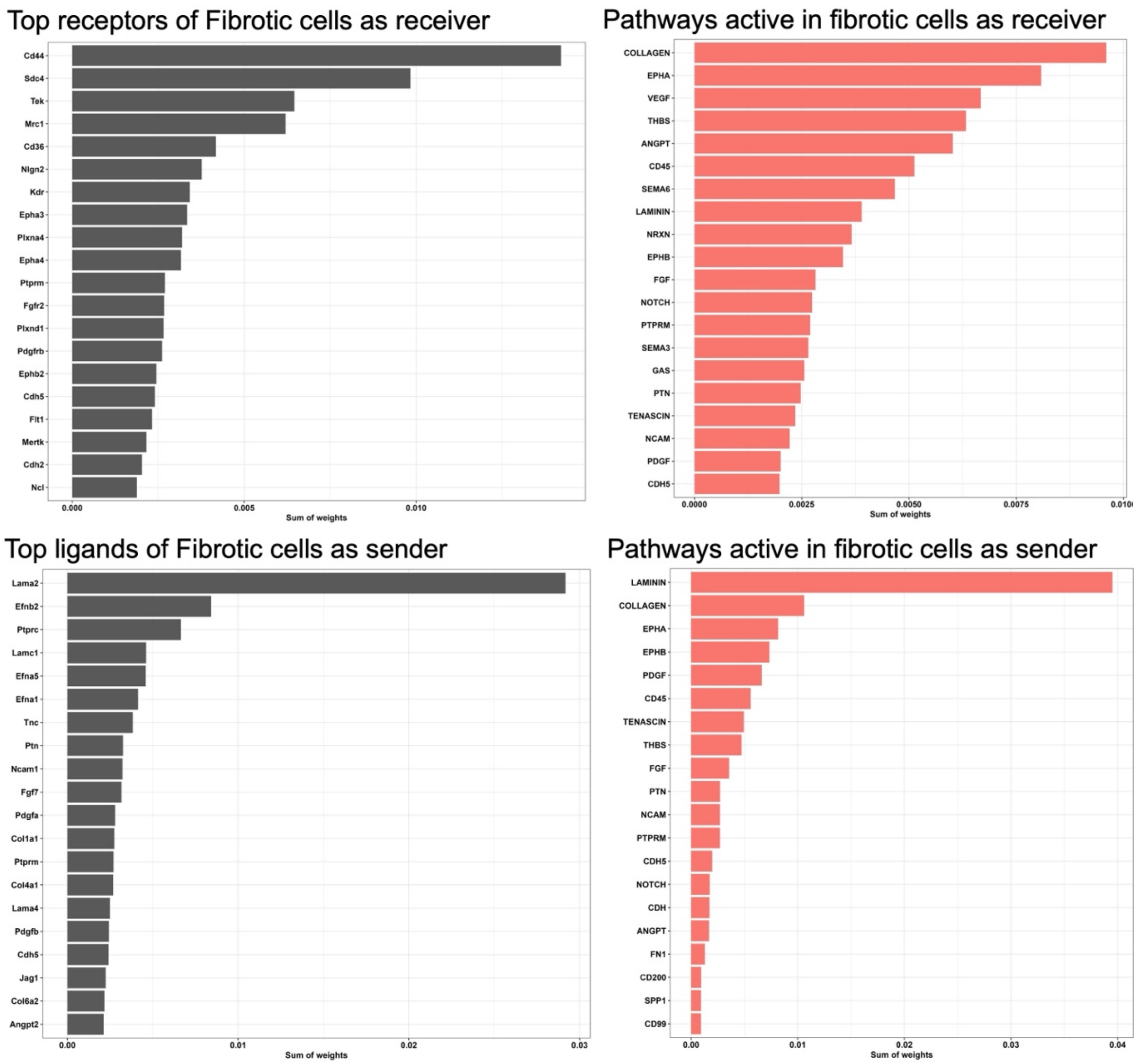


**Supplementary Figure S4.** Receptor, ligand, and pathway interaction profiles associated with fibrotic cells in CellChat analysis.

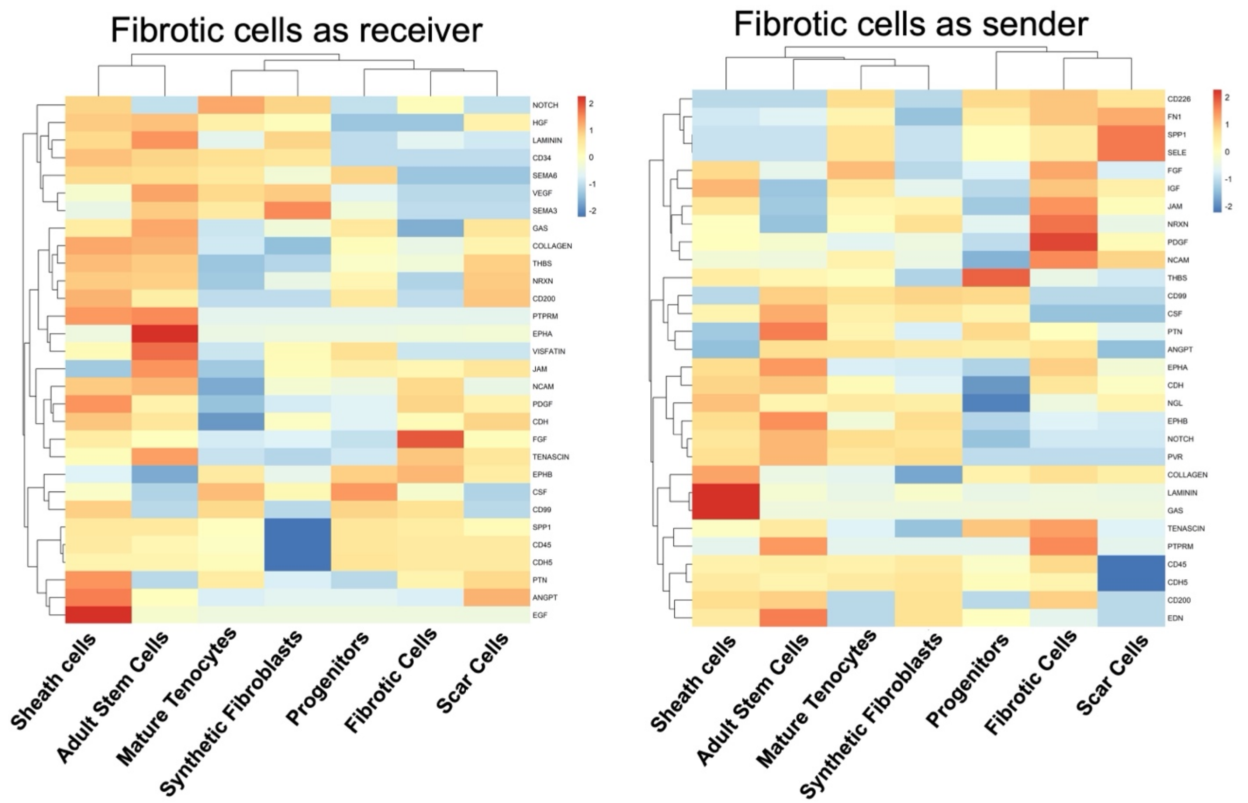


**Supplementary Figure S5.** Signaling pathway activity associated with fibrotic cells as receivers and senders

### Supplemental Methods

#### Tendon injury

Following injury, the skin was closed with bioresorbable suture. Rats were placed in a solitary recovery cage with ad libitum access to food and water.  Buprenorphine XR 65mg/kg was administered after operation. At each of three pre-specified post-injury time points, 14 rats (total n = 42) were euthanized by $CO_2$ inhalation immediately followed by lung puncture to ensure death for single nuclei RNA-sequencing study, in accordance with institutional IACUC approval (No. 008586). Based on an a priori power analysis, 13 animals per group per time point were required to achieve 80% power at a two-sided $\alpha$ of 0.05 for the primary outcome measures. Therefore, one additional animal was included per group/time point to account for potential attrition. Tendon tissues were promptly harvested for nuclei isolation. Following biobehavioral analysis, total 28 rats (n=14/sample) were euthanized for biomechanical, histology and immunofluorescence analyses. The contralateral Achilles tendons served as control group for all studies.

#### Gait Analysis

Stride length was measured as the distance between the proximal points of successive thenar pads from the same paw. Stride width represented the perpendicular distance between the midpoints of opposite strides. The paw angle for external rotation was determined relative to the rat's forward axis by drawing a circle through the proximal points of the second to fourth metatarsals, a line from the circle's center through the thenar pad to the midline defined the measured angle. Individual right paw angles were converted to adjusted values using a pre-established transformation, and replicate measurements within each timepoint were averaged to obtain one mean right and one mean left value per rat per timepoint. The paw width was defined as the distance between the distal ends of the first and fifth phalanges. The normalization was performed within each animal by dividing each week-specific mean by the corresponding baseline mean for that paw. The functional gait analysis was performed using Achilles Functional Index (AFI) as previously described[1]. Data was blindly calculated to prevent observer bias by MB, LZ and MO. The animal log was kept by AEP and JS.

#### Biomechanical Analysis

After collecting the data from the uniaxial mechanical testing device, the fold change in CSA was calculated by dividing the injured values to uninjured values obtained for each AT and YT sample. Afterwards, each tendon was mounted onto dry ice-frozen needle holders, specifically the uninjured section of the tendon at tendon-muscle junction and placed above the calcaneus, ensuring secure fixation without slippage during testing. Tensile testing was performed until break at a speed of 1 mm/min using uniaxial mechanical testing device (Hydraulic mechanical testing system 370.02 Bionix, MTS Systems Corp., Eden Prairie, MN). Tendons were failed the mid-substance. The load determined at tendon rupture was denoted as failure load. Using the CSA and failure load, tensile strengths (MPa) and elastic moduli (MPa) were calculated. Biomechanical outcomes were normalized to the contralateral uninjured tendon for each animal to quantify injury-induced changes within AT and YT groups. The contralateral tendons served as uninjured controls.

**Histology and immunofluorescence**

The tendons were placed in 4% paraformaldehyde overnight. Then, the samples were dehydrated by running them through graded ethanol series and finally they were embedded in paraffin. Five micron-thick sections were obtained, re-hydrated and stained with hematoxylin and eosin (H&E) for morphological feature evaluation. For immunofluorescence (IF), re-hydrated tendon sections were treated with Proteinase K to retrieve antigens for 20 min at room temperature. Nonspecific antigens were blocked by applying blocking agent (Serum-free Protein Block, Doku). The primary antibodies listed in Supplemental Table 1 were applied for 1 h at room temperature. Then, tendon sections were rinsed thoroughly with 2% bovine serum albumin in PBS three times and treated with secondary antibodies for 30 min at room temperature. After rinsing with 2% BSA several times, the sections were mounted on cover slips using antifade mounting medium (Fluoromount-G with Dapi, Invitrogen) for confocal laser scanning microscopy analysis (Nikon AX/AXR Confocal Microscope System, Japan).

**Supplemental Table 1.** Antibodies and dyes used in IF study.

<table>
<tr><th>Staining sets</th><th>Primary Antibodies</th><th>Host species</th><th>Secondary Antibodies</th><th>Host species</th></tr>
<tr><td rowspan="3">Set 1</td><td>Scx</td><td>Rabbit (PA5-115874, Invitrogen)</td><td>Anti-rabbit Cy2</td><td rowspan="6">Donkey (AffiniPure IgG, Jackson ImmunoResearch Laboratories)</td></tr>
<tr><td>Fabp4</td><td>Goat (AF3150-SP, R&D)</td><td>Anti-goat Cy3</td></tr>
<tr><td>Gas6</td><td>Mouse (SC-376087, Santa Cruz Biotechnology)</td><td>Anti-mouse Cy5</td></tr>
<tr><td rowspan="3">Set 2</td><td>Tnmd</td><td>Rabbit (HPA055634, Sigma Aldrich)</td><td>Anti-rabbit Cy2</td></tr>
<tr><td>Tnc</td><td>Goat (MBS421644, MybioSource)</td><td>Anti-goat Cy3</td></tr>
<tr><td>Lrp6</td><td>Mouse (SC-25317, Santa Cruz Biotechnology)</td><td>Anti-mouse Cy5</td></tr>
</table>

**Single Nuclei RNA Sequencing**

Single nuclei suspensions were then stained with ViaStain AOPI Staining Solution (Nexcelom Bioscience, Lawrence, MA) and imaged on an EVOS M7000 Imaging System (Thermo Fisher Scientific, Waltham, CA) to determine cell suspension quality and cell viability. Nuclei suspensions were partitioned (1500 nuclei/mL target capture) into Gel Beads-in-emulsion (GEMs) on the Chromium X (10x Genomics, Pleasanton, CA) per the Chromium GEM-X Single Cell 3' v4 Gene Expression kit user guide in Cedars Sinai Medical Center Applied Genomics and Translational Medicine Core (CSMC AGTC). Following reverse transcription and cDNA amplification, single cell libraries were prepared on the Chromium Connect (10x Genomics) using the Automated Gene Expression Library Construction kit (10x Genomics). Barcoded libraries were quantified by quantitative PCR using the Collibri Library Quantification Kit (Thermo Fisher Scientific) and library size was measured via the 4200 TapeStation (Agilent Technologies, Santa Clara, CA). Libraries were sequenced on a NovaSeq X Plus (Illumina, San Diego, CA) with sequencing configuration 28x10x10x90 bp, and a sequencing depth of 20K reads/nuclei. After quality control, low feature nuclei were removed.

Seven libraries were profiled and yielded 60479 nuclei before QC steps: Uninjured AT 10176, AT_w1 13150, AT_w2 5478, AT_w4 8116, YT_w1 12551, YT_w2 4324, YT_w4 6684. 10x matrices were imported with Seurat and converted to Seurat objects (min.cells = 3, min.features = 200). Mitochondrial content was computed with "PercentageFeatureSet" using the pattern "^MT-|^Mt-|^mt-". Cells were retained if percent.mt < 10% and 200–8,000 genes were detected. Each sample was normalized and variance-stabilized with "SCTransform" (method = "vst", defaults). Sample identities were harmonized via an ordered "orig.ident" factor with levels: Uninjured, AT_w1/2/4 and YT_w1/2/4. "SCTransformed" objects were integrated using "SelectIntegrationFeatures" (3,000 features), "PrepSCTIntegration", "FindIntegrationAnchors" (normalization.method = "SCT"), and "IntegrateData" (normalization.method = "SCT"). The integrated object was reduced by PCA, embedded by UMAP (dims = 1:20), and graph-built with "FindNeighbors" (dims = 1:20). Cluster markers were identified with "FindAllMarkers" (defaults; only.pos = TRUE, min.pct = 0.25, logfc.threshold = 0.25). Cells annotated as CTC were subset and reprocessed with "SCTransform", PCA, UMAP (dims = 1:20), and "FindNeighbors" and "FindAllMarkers" and visualized by heatmaps and dot plots. Curated gene sets were scored per cell with "AddModuleScore" on the SCT assay and clamped to [−2, 2] for comparability across panels. Ingenuity Pathway Analysis (QIAGEN IPA) was run on cluster marker lists. Excel outputs were parsed in R and summarized as top-10 positive/negative z-score pathways ordered by −log10(p); figures were produced for the whole-dataset synthetic fibroblasts and fibrotic cells.

**Slingshot trajectory analysis.** To resolve reparative versus fibrotic paths within CTCs, we performed unsupervised trajectory inference with Slingshot via "SingleCellExperiment"[2]. Lineages were inferred on the UMAP embedding using cluster labels, principal curves were overlaid with directionality arrows.

**CellChat analysis.** Ligand–receptor communication was inferred with CellChat from a Seurat object of the CTCs, with cells grouped by clusters. The full CellChat rat database was used to compute pathway-level communication probabilities, which were aggregated by sender (source cluster) and receiver (target cluster). Analyses targeted fibrosis-centric interactions: 1) interaction-strength heatmaps for partners → fibrotic cells and fibrotic cells → partners, and 2) overview sender/receiver heatmaps summarizing pathway activity across clusters. These visualizations highlight dominant incoming signals to the fibrotic population and its outgoing influence on neighboring compartments.

**Statistical analysis**

The significant differences were denoted as *$p<0.05$, **$p<0.01$, ***$p<0.001$ and ****$p<0.0001$. Data points over two standard deviations were excluded from analysis as outliers. For gait analysis, 2-way analysis of variance (ANOVA) analysis was performed. For multiple comparisons, appropriate post hoc tests were used. The sample sizes for animal studies were calculated according to our previous work.
